# Viral delivery of recombinases to activate heritable genetic switches in plants

**DOI:** 10.1101/2024.03.03.583219

**Authors:** James C. Chamness, Jon P. Cody, Anna J. Cruz, Daniel F. Voytas

## Abstract

Viral vectors provide an increasingly versatile platform for transformation-free reagent delivery to plants. RNA viral vectors can be used to induce gene silencing, overexpress proteins, or introduce gene editing reagents, but they are often constrained by carrying capacity or restricted tropism in germline cells. Site-specific recombinases that catalyze precise genetic rearrangements are powerful tools for genome engineering that vary in size and, potentially, efficacy in plants. In this work, we show that viral vectors based on *Tobacco rattle virus* (TRV) deliver and stably express four recombinases ranging in size from ∼0.6kb to ∼1.5kb, and achieve simultaneous marker removal and reporter activation through targeted excision in transgenic *Nicotiana benthamiana* target lines. TRV vectors with Cre, FLP, CinH, and Integrase13 efficiently mediated recombination in infected somatic tissue, and also led to heritable modifications at high frequency. An excision-activated Ruby reporter enabled simple and high-resolution tracing of infected cell lineages, without the need for molecular genotyping. Together, our experiments broaden the scope of viral recombinase delivery, and offer insights into infection dynamics that may be useful in the development of future viral vectors.

## Introduction

Genome engineering in plants is limited by challenges in reagent delivery. Due to the difficulty of stable transformation, numerous plant viruses have been repurposed as systemically mobile vectors for applications including virus-induced gene silencing (VIGS), virus-mediated protein overexpression (VOX), and virus-induced gene editing (VIGE). Benefits of viral delivery include reduced researcher labor and faster timescales versus sterile culture, and high-level reagent expression through viral replication. Drawbacks include narrow host ranges, deleterious symptoms in infected plants, and tissue tropisms that limit reagent access to target cells, particularly meristems. Additionally, many viruses have limited capacity to add exogenous sequences, above which mobility is attenuated or selective pressure results in vector collapse and loss of cargo.

Recent progress has been made developing a first generation of VIGE vectors. Two reports used viral vectors based on *Sonchus yellow net rhabdovirus* (SYNV) and *Potato virus X* (PVX) to deliver both Cas9 and single guide RNA sequences (sgRNAs) to wild-type plants, but seed transmission of events required regeneration of edited tissue through sterile culture^1,2^. In culture-free approaches, two viruses capable of meristem invasion, *Tobacco rattle virus* (TRV) and *Barley stripe mosaic virus* (BSMV), have been used to deliver sgRNAs to Nicotiana, Arabidopsis, and wheat^3–7^. These approaches rely on transgenic lines, as TRV and BSMV cannot carry the longer coding sequences for large, Cas-derived proteins. The range of heritable modifications achieved with viral delivery of RNA-guided editing tools has thus far been limited to mutagenesis or base editing.

Another significant class of genome engineering reagents are site-specific recombinases (SSRs). With different placements and orientation of their cognate recognition sites, SSRs can mediate recombination reactions including excision, integration, inversion, and cassette exchange. The coding sequences for many SSRs are considerably shorter than most Cas proteins, and several previous reports have demonstrated viral delivery of Cre recombinase (∼1kb) to transgenic *Nicotiana benthamiana, Nicotiana tobaccum*, and potato reporter lines, using either PVX or *Tobacco mosaic virus* (TMV). Most of these reports required a sterile culture step to regenerate fixed recombination events from infected tissue and, for PVX, to eliminate infectious virus^8–10^; one study claimed germline transmission of recombination in *N. benthamiana* using a PVX-Cre vector, but at low frequency among infected plants, and data was not presented to confirm absence of the virus among the progeny^11^. These results are consistent with the understanding that both PVX and TMV are generally excluded from meristems^12^, a likely prerequisite for efficient germline transmission.

TRV is an attractive vector for viral delivery with a broad host range, mild symptoms, and ability to transiently invade meristems without passing to progeny through seed^12,13^. In this study, we developed *N. benthamiana* target lines with a recombination-activated Ruby reporter^14^ for four different recombinases, and infected T_1_ plants from each line with a corresponding TRV recombinase vector. The viral vectors were sufficiently stable, even with the largest recombinase, to produce robust recombination through infection, including heritable events among virus-free progeny. Our work demonstrates an improved method for viral delivery of recombinases, and illustrates a highly practical strategy for studying the infection patterns of viral vectors.

## Results

### Recombinase switch design for reporter activation by viral infection

We designed a series of genetic switch constructs to function in transgenic plant lines as targets for viral infection with different recombinases. In the construct archetype, a *2×35S* promoter drives the selectable marker *NptII*, followed by a promoter-less Ruby reporter. Recombinase recognition sites are placed immediately upstream of both coding sequences in the same orientation, so that recombination results in excision of the *NptII* selectable marker and activation of the Ruby reporter behind the same promoter (Figure 1A). We first built two constructs and generated transgenic *N. benthamiana* lines implementing this scheme with different pairs of *lox* sites for Cre recombinase, modified to favor unidirectional recombination: pJPC259, using *lox71* and *lox66* ^15^, and pJPC263, using *loxP1L* and *loxP1R*^16^. Among previously developed *N. benthamiana* transgenic lines with a fixed *2×35S::Ruby* cassette, we observed highly variable intensity of the red pigmentation phenotype, likely attributable to event-to-event variation in expression levels of the triple gene block encoding the betalain biosynthetic pathway. To identify target line events with strong Ruby expression, we screened T_1_ seed using the Fast-TrACC protocol for Agroinfection of germinating seedlings^17^ to deliver Cre recombinase on TRV. Based on this screen, we selected one event per construct, 259-12 and 263-8, to proceed with new infections of soil-grown T_1_ plants.

**Figure 1:**
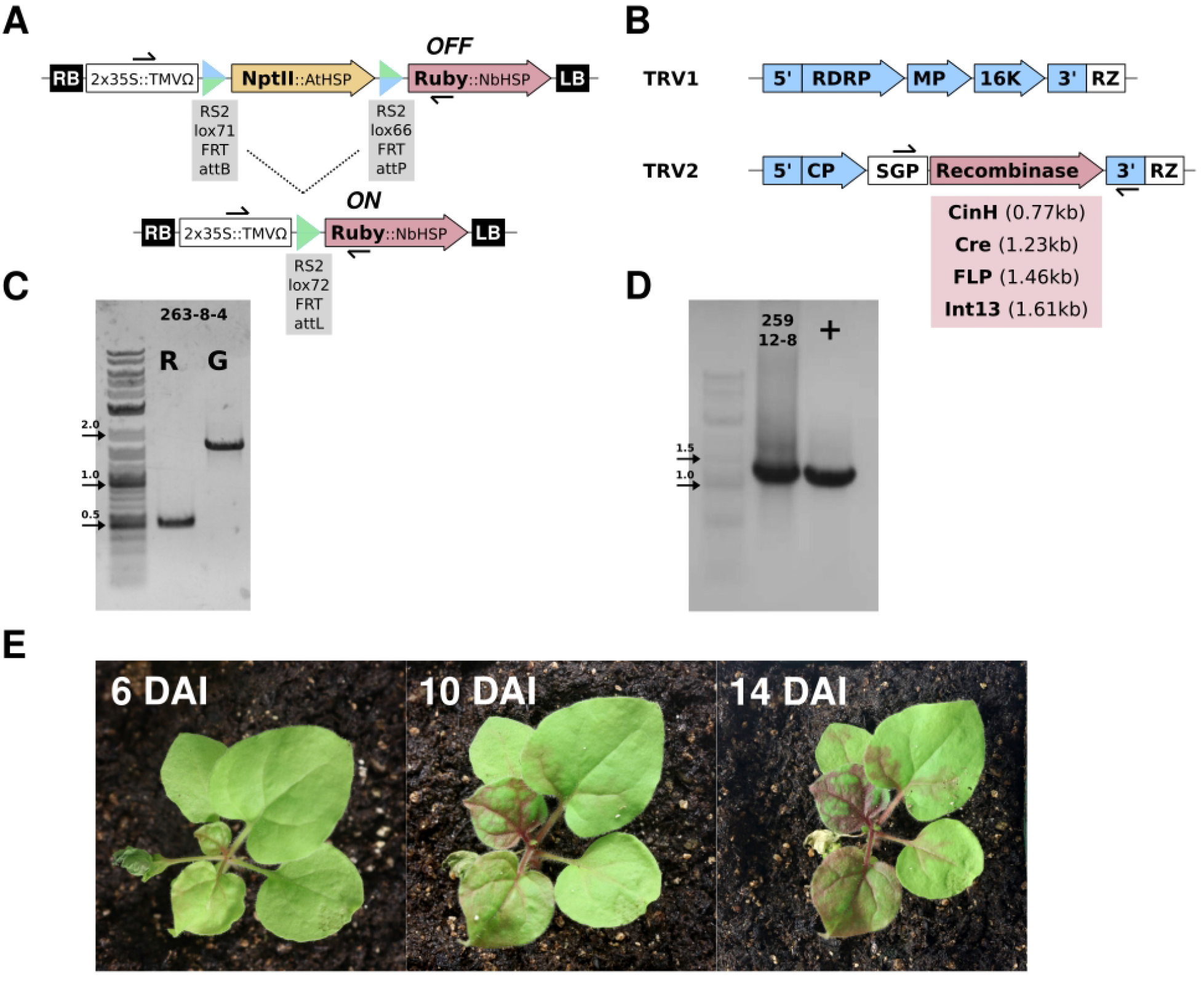
NptII/Ruby switch actuated by recombinase delivery via TRV. **1A:** Vector schematic for transgenic target lines illustrating pre- and post-recombination forms. Black arrows indicate primer binding sites for diagnostic PCR. **1B:** Vector schematic for recombinase delivery on TRV, including unmodified TRV1 and TRV2 with the PeBV subgenomic promoter (SGP) driving the recombinase. Both subgenomes were delivered on T-DNA vectors with a 2×35S promoter (not indicated) and 3’ ribozyme site (RZ). Black arrows indicate primer binding sites for RT-PCR. **1C:** Diagnostic PCR for TRV-Cre-mediated excision in infected target line: lane R indicates tissue sampled from Ruby-positive red tissue, and lane G indicates tissue sampled from green tissue. Predicted full-length and recombination bands are 1683bp and 511bp, respectively. **1D:** RT-PCR for TRV-Cre: + indicates plasmid control, and labeled lane indicates RNA sampled from target line 292 days after infection. **1E:** Timelapse of emerging Ruby phenotype from 6 to 14 days after infection.

TRV-Cre infection of T_1_ plants was initiated by co-infiltration of TRV1 and a modified TRV2, featuring Cre downstream of the TRV coat protein and driven by the *Pea early browning virus* subgenomic promoter (Figure 1B). Soon after infiltration, red pigmentation began to appear in the stem above the infiltrated leaf, spreading through the vasculature and into leaves (Figure 1E). To confirm the phenotype was attributable to recombination, we sampled genomic DNA from green and red tissue within one of the infected plants, 263-8-4, and performed PCR with primers flanking the recombination junctions. The green tissue produced the expected full-length band, while the red tissue produced the expected shorter band. Furthermore, the absence of any full-length band in the red sample indicated a high level of recombination within the infected tissue. Extraction and sequencing of both bands produced perfect alignments to the source plasmid and the predicted *loxP1LR* recombination sequence, respectively (Figure S1). Ruby signal continued to spread over weeks and even months of plant development, suggesting continued TRV infection and stability of the recombinase insert. To confirm systemic movement and stability of the TRV-Cre vector directly, we extracted RNA from red, upper leaves of an infected plant and performed RT-PCR with primers flanking the Cre coding sequence on TRV2. The insert was very stable; remarkably, one RNA sample collected 7.5 months after infection still produced a strong, full-length band (Figure 1D). Together, these data validated the Ruby switch as an effective system for scoring viral infection-mediated recombination with Cre.

### High-resolution tracing of TRV infection lineage with Ruby

During the course of development, we observed variation in Ruby patterning among the infected plants which was nonetheless consistent with expectations for viral movement. Most plant viruses rely on the vasculature for long-distance transport, and plasmodesmata for localized movement into and between mesophyll and epidermal cells^18^. Outside of the infiltrated leaf, Ruby signal in the TRV-Cre infected plants first emerged in the main shoot, then spread into axillaries and leaves in a fashion traceable down to individual leaf veins (Figure 2A, 2C). Wild-type TRV is a nematode-transmitted virus, and previous studies have shown that TRV2 vectors expressing GFP exhibit tropism to the roots, even with removal of the 2b protein required for nematode transmission^19,20^. Our TRV-Cre vector, which also omits the 2b protein, similarly gave rise to Ruby signal in the roots of examined plants (Figure 2B). Most critically for achieving heritable modifications, we also observed Ruby signal in reproductive organs once the infected plants began to flower, including red-streaked or fully-red pods, sepals and petals (Figure 2D). In multiple tissues, the intensity of Ruby signal increased with time, likely reflecting both increased penetrance of viral infection and viral-independent accumulation of betalain in previously infected recombinant cells.

**Figure 2:**
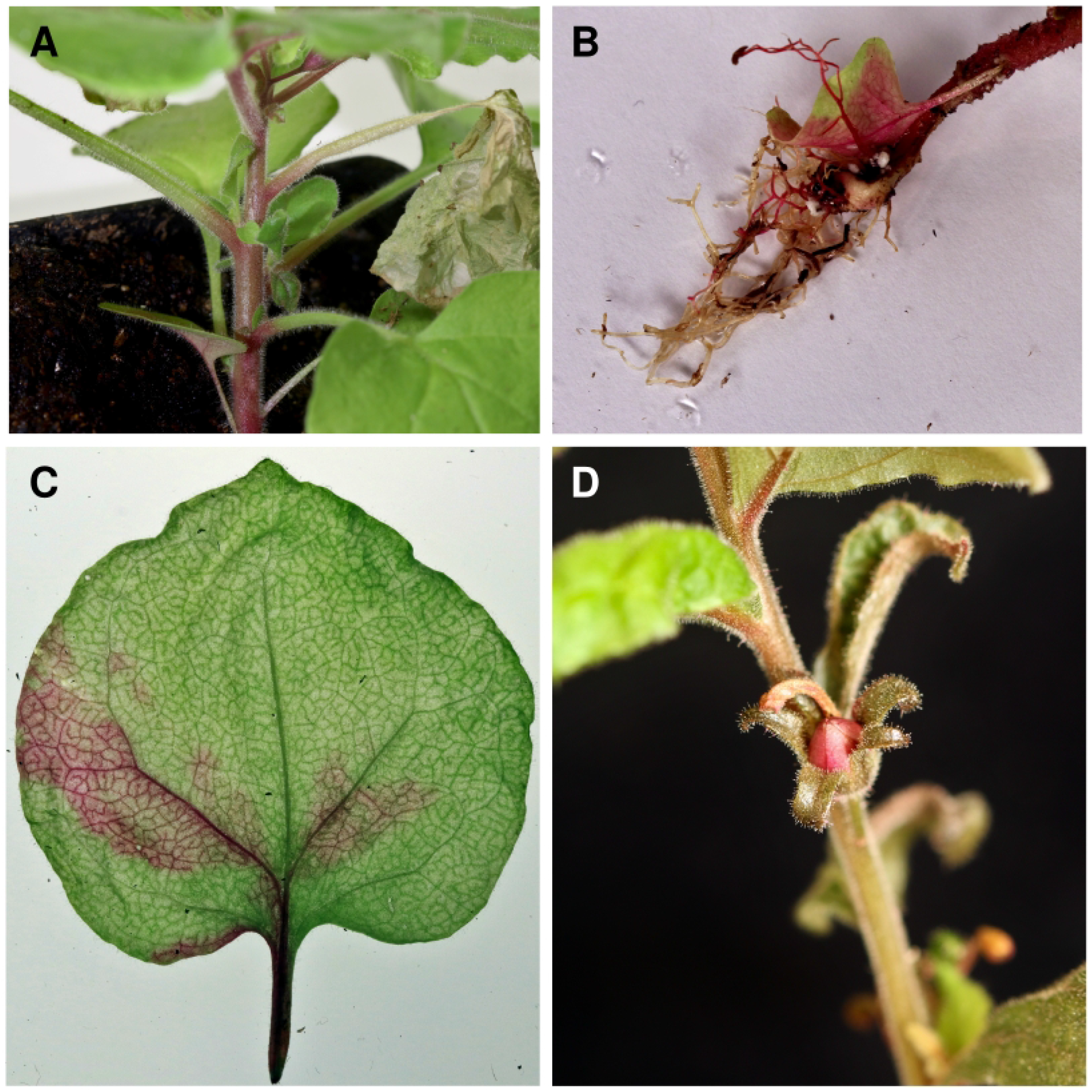
Tracing TRV infection with Ruby reporter. **2A:** Initial signal emergence in the stem is consistent with vasculature-mediated systemic transport of TRV. **2B:** Tropism to roots is consistent with nematode transmission for the wild-type virus. **2C:** Mesophyll and epidermal cell infection initiates from the vasculature. **2D:** Ruby signal in floral organs indicates infection and recombination in germline cells.

### Heritable recombination events from TRV-Cre infection

A primary goal for VIGE experiments, and for recombinase applications such as marker removal, is to generate heritable modifications. The presence of Ruby-positive flowers suggested TRV-Cre successfully invaded germline cells, which we sought to confirm by phenotype and molecular genotype among T_2_ progeny from the infected plants. One barrier we encountered was low viability and fertility among the infected T_1_ parents, partially accountable to our experimental design. For each event, we initially infected 5 plants grown at each of two different growth regimes: 21°C and 26°C. The lower temperature is more permissive, leading to virulent TRV infection and more severe symptoms in *N. benthamiana*. Each set of 5 plants was infiltrated at a range of developmental stages to explore the effect of plant age on infection efficiency. Across all 20 plants, 10 showed at least some Ruby signal, but many did not survive to adulthood: this was especially true for the groups grown at 21°C, and those plants infiltrated very early in development (Figure S2). These T_1_ plants were also still segregating for the T-DNA insertion(s), and we did not perform any genotyping or selection prior to infection, so some of the infected plants may have been wild-type. In total, we recovered and screened seed from 3 of the 10 Ruby-positive plants.

The first plant, 259-12-8, was grown at 21°C and infected early in development, yet grew normally and produced many viable seed despite widespread TRV-Cre infection (Figure 3A). For this plant, we collected and screened progeny pod-by-pod. The first pod produced 11 seed: 10 Ruby positives, and 1 entirely green (Figure 3C, 3D). Of the 10 Ruby positive plants, 9 were PCR positive for recombination. The single green plant (#11) produced only the full-length band in PCR (Figure 3B). From pod 2, all 46 progeny were Ruby positive, and 45 of these were PCR-positive (Figure S3A). From across pods 3 and 4, 7 of 11 sampled progeny were PCR-positive (Figure S3B). The second plant, 259-12-1, was grown at 21°C, and produced a high fraction of viable progeny which we screened as a bulk collection across seed pods. From 77 seedlings, 69 were Ruby positive (Figure 3E), and 13 of 14 subjected to PCR were positive with a single band for recombination (Figure S3C). The third plant, 263-8-4, displayed severe stunting (Figure S3D), and only 8 of 52 collected seed germinated. All 8 were positive for recombination, but died before a phenotype was clearly discernible (Figure S3E). Despite the low rate of plant viability, these data indicate frequent vertical transmission of recombination events among infected plants, independent of pod position, and with clear co-segregation of molecular and phenotypic markers. Lastly, although TRV is considered non-seed transmissable, we decided to screen four progeny, from 259-12-8 pod 1, via RT-PCR for the presence of TRV-Cre: as expected, none of the 4 plants were positive (Figure 3F).

**Figure 3:**
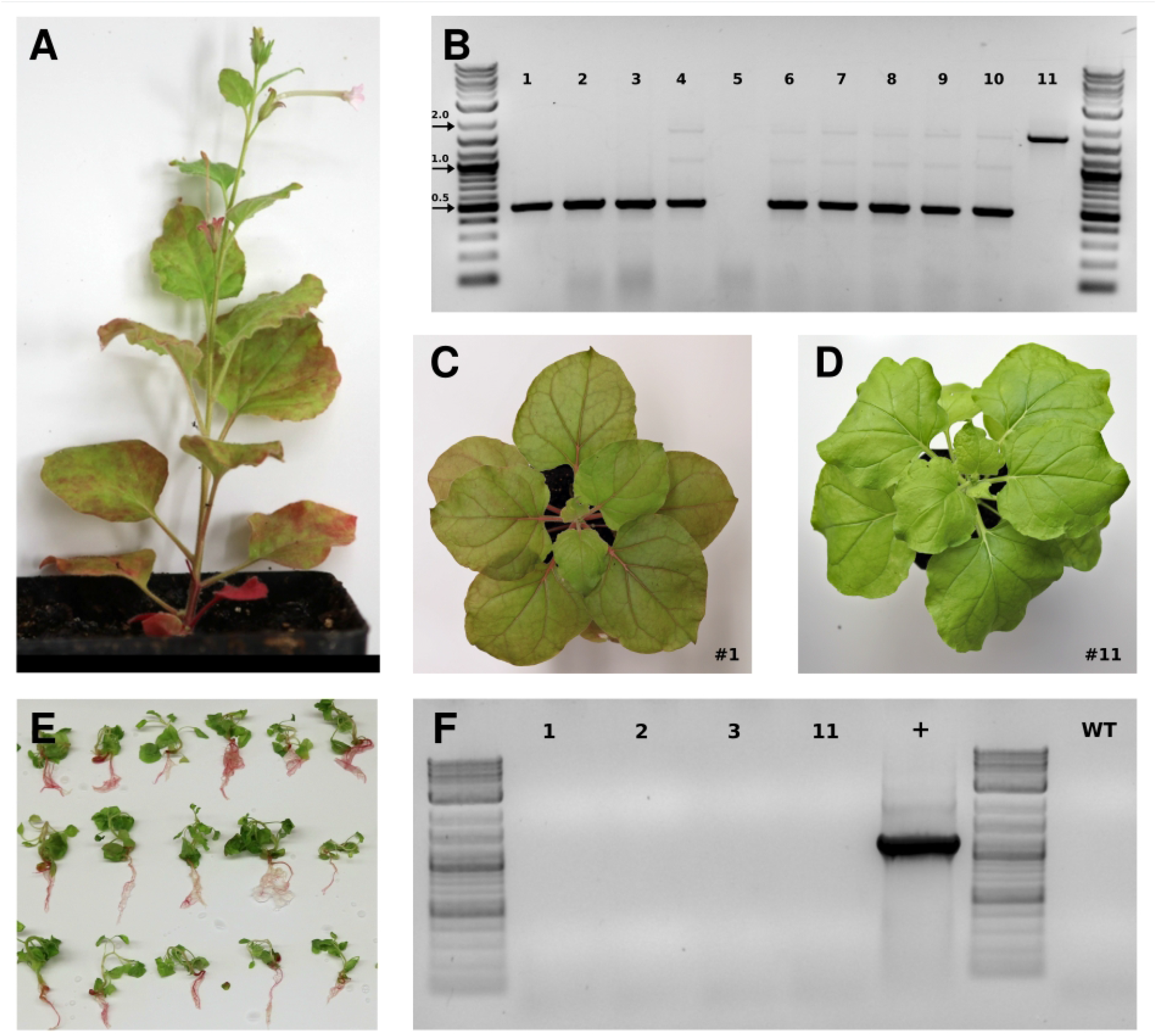
TRV-Cre recombination events are heritable at high frequency. **3A:** Infected parent plant 259-12-8 at flowering age, showing moderate Ruby phenotype. **3B:** Diagnostic PCR for 11 progeny from the first seed pod of 259-12-8. **3C:** Progeny plant #1 from the set in 3B, with moderate Ruby phenotype. **3D:** Progeny plant #11 from the set in 3B, with no Ruby phenotype. **3E:** Subset of seedlings germinated from bulked seed from plant 259-12-1, with Ruby visible in the roots of nearly every plant. **3F:** RT-PCR for TRV-Cre in subset of progeny from 3B: + indicates plasmid control, and WT indicates RNA from a wild-type plant.

### Phenotypic defects among TRV-Cre infected plants

Low plant viability among the Cre target lines could be due to viral infection and strong symptoms under permissive conditions, as discussed above. However, another contributing factor may be the recombinase itself. Constitutive and/or high-level Cre expression, particularly in early development, has been associated with deleterious effects, including chromosomal rearrangements in mammalian cells, and phenotypic abnormalities in plants. In mammalian studies, these effects depend on the Cre endonuclease function, and likely arise from recombinase action on cryptic *lox* cognates present in the host genome^21–23^. To disaggregate T-DNA insertion location and high-level betalain production as other potential effectors, we infected six wild type plants with TRV-Cre: all these plants, as well as several infected transgenic lines that never produced Ruby signal, exhibited phenotypic defects, including stunting, misshapen leaves, and loss of apical dominance (Figure S4).

### TRV delivery of FLP, CinH, and Integrase13

While Cre is the most widely used SSR, others have also been shown to function in plants. We reasoned that, with different recognition sites, alternative recombinases may sidestep the phenotypic penalties observed with TRV-Cre. SSRs also differ by the classes of reactions they catalyze, and in the sequence and activity of their recognition sites pre- and post-recombination. Marker removal can be accomplished with any excision reaction, but more advanced applications such as gene stacking would also benefit from a broader toolbox of functional recombinases. We thus selected three additional SSRs to conduct analogous experiments: FLPe, another bidirectional tyrosine recombinase, and CinH, a small, unidirectional serine recombinase, both of which have previously been used for marker removal in transgenic plants^24,25^; and Integrase13, a large serine integrase, for which the only available plant data is from protoplasts^26,27^.

We generated transgenic *N. benthamiana* lines with similar target constructs, substituting recognition site pairs for each new SSR: pJPC430, with identical FRT sites for FLP; pJPC432, with identical RS2 sites for CinH; and pJPC210, with attB/attP sites for Integrase13 (Figure 1A). Unlike the Cre target lines, we did not pre-screen events by Fast-TrACC, but proceeded directly to TRV infections of four soil-grown T_1_ plants per event. These plants were grown at 25°C and, where possible based on asynchronous germination, infiltrated past the fourth-true-leaf stage. Between two to three weeks after infection, Ruby signal began to emerge in plants from events across each construct. As with the Cre-*lox* lines, there was considerable variation in intensity of the Ruby phenotype. We focused on a subset of plants from 6 different events with high expression: plants 7-1, 10-1, 12-1, and 13-2 with Integrase13; plant 5-1 with FLP; and plants 1-1, 1-2, 1-3, and 1-4 with CinH. While all four infected plants from CinH event #1 produced signal, these plants were slow to germinate and younger than the others at the point of infiltration, which may have affected viability. Two died soon after infiltration, and another was abnormally small, leading us to genotype only plant 1-1.

At 24 days after infection, we collected tissue to genotype the six plants described above with strong signal. Using RNA extractions, we conducted RT-PCR with the same primers to probe stability of the recombinase inserts on TRV (Figure 4A). The smallest sequence, CinH, was perfectly stable; the largest sequence, Integrase13, exhibited collapse at variable frequency, but did produce a full-length band in every plant; no full-length band was detected for FLP. We suspect this was due to sampling from lower leaves that were either never infected or where the virus collapsed - an RNA sample taken later from upper leaves that turned red was positive for the full-length virus, with a modest degree of collapse (Figure 4B). Using genomic DNA extractions, we conducted a PCR diagnostic, where every sample was positive for a lower band indicative of recombination (Figure 4C). We extracted and cloned these bands, then sequenced several independent clones. Interestingly, while every clone sequence indicated that excision took place, some clones showed unanticipated small deletions at the recombination junction, either within or flanking the expected final recognition site sequence (Figure S5). A mixed population of recombination excision products, while unexpected, is consistent with the faint double banding observed in the original PCR gel.

**Figure 4:**
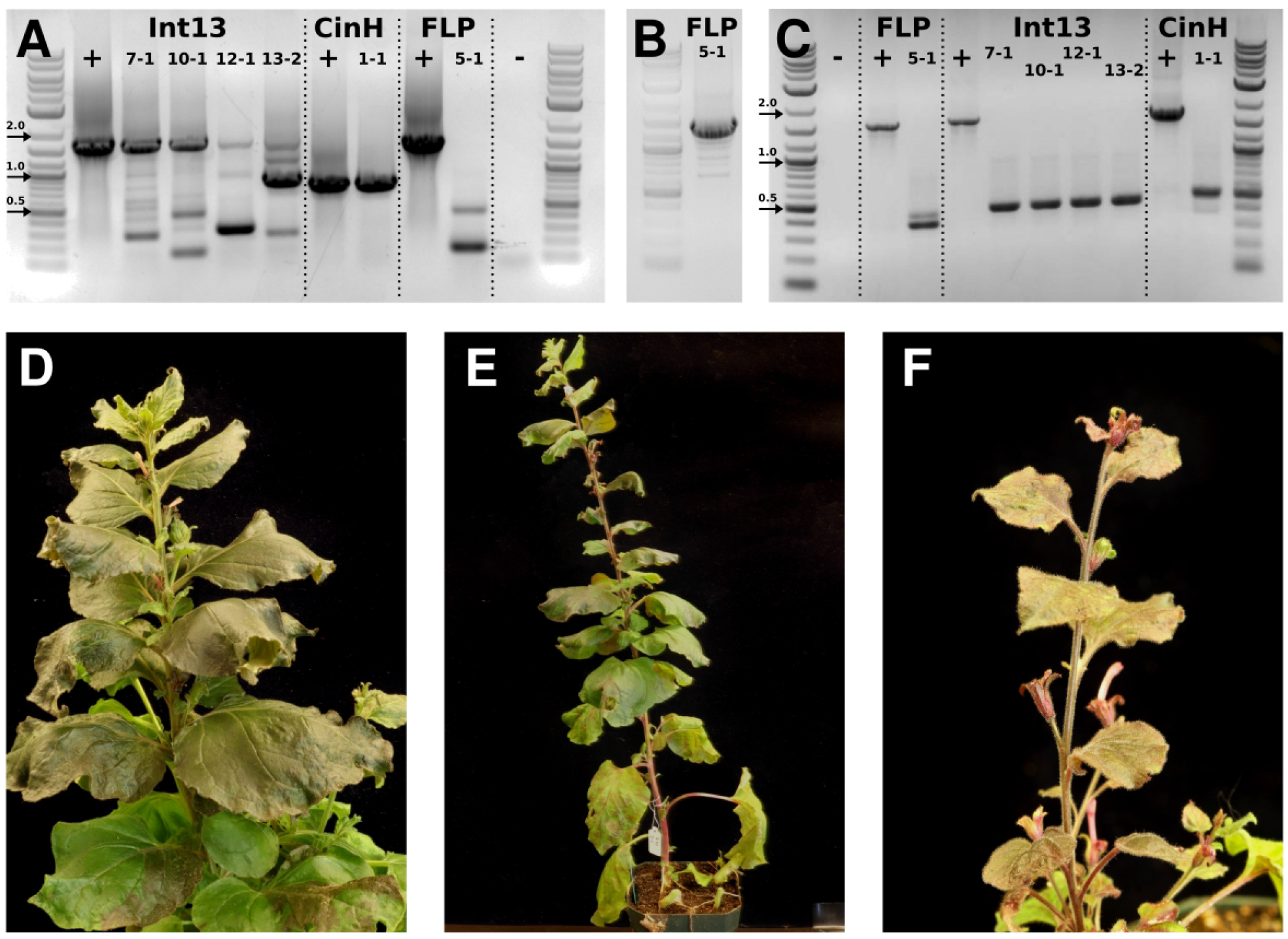
Viral delivery of additional recombinases. **4A:** RT-PCR for different viral recombinases in infected plants: + indicates plasmid control, - indicates water negative control, and labels indicate distinct event/plant samples. Full-length bands are 1462bp for FLP, 1608bp for Int13, and 774bp for CinH. **4B:** RT-PCR for FLP in upper leaf tissue. **4C:** Diagnostic PCR for recombination in Ruby-positive tissue of infected plants. Full-length and recombination bands are 1662/477bp for Int13, 1770/532bp for CinH, and 1600/447bp for FLP. **4D-F:** Infection activates Ruby expression in infected plants, including flowers, for all 3 recombinases. **4D:** TRV-FLP infection in plant 432-5-1. **4E:** TRV-Int13 infection in plant 210-7-1. **4F:** TRV-CinH infection in plant 430-1-2.

For the majority of plants, Ruby signal spread throughout months of development, indicating continued infection. For FLP, plant 5-1 remained green in most lower leaves, but the entire upper portion of the plant turned red (Figure 4D). For Integrase13, plant 10-1 never produced Ruby signal beyond the stem above the infiltrated leaf, but plants 7-1, 12-1, and 13-2 eventually turned red through nearly the entire vasculature (Figure 4E). For CinH, plant 1-1 died prior to flowering, but the surviving plant 1-2 turned entirely red; the small stature of this plant motivated us to preserve leaf integrity, rather than sample punches for genotyping (Figure 4F). With the exception of the small plants from CinH event #1, none of the infected plants exhibited the developmental abnormalities previously associated with Cre expression. For all plants where visible Ruby signal indicated widespread infection, we observed at least some signal in reproductive organs when those plants reached flowering.

In addition to the events with strong systemic signal, we observed different Ruby phenotypes among infected plants from other events. This may result in part from variegated infection by TRV, but we suspect most observations reflect expression-level variation between events, as with the Cre-*lox* lines assessed in Fast-TrACC. In some cases, faint signal took longer to accumulate to visible levels, or was only visible in a subset of tissues (typically vasculature). Additional plants remained entirely green throughout development until they produced pale pink flowers, likely the result of stable infection in events where Ruby expression was too low to visualize in green tissue (Figure S6). Together, our results show that all three additional recombinases are sufficiently stable on TRV to facilitate recombination and Ruby reporter activation in the corresponding target lines.

### Screening for heritable recombination events from TRV-FLP, CinH, and Integrase13 infections

Given efficient vertical transmission of recombination events from the TRV-Cre infected plants, we next determined if the other TRV recombinase vectors could produce heritable recombination events. For CinH, only one of the 4 infected plants from the high expression line, 430-1-2, survived to flowering. Seed set from this plant was poor, yielding only 7 seeds from a bulk across all pods. To maximize viability, we germinated these in vitro, without selection, resulting in a mix of red and green seedlings. Given the small sample size and uncertain zygosity of the T-DNA in the infected parent, we screened each seedling by PCR for both a genomic control locus, *AGAMOUS*, and the T-DNA recombination probe, so that empty lanes for the latter could confidently be attributed to null segregants (rather than gDNA extraction or PCR failure). All 6 seedlings produced the genomic control band, and 4 produced a band for the T-DNA probe: 1 full-length, and 3 with the recombination band, a heritable recombination rate of 75% among transgenic progeny (Figure 5A). Given the unexpected deletions observed within sequencing of some CinH recombination bands in the infected generation, we also cloned and sequenced the diagnostic amplicon from one of the recombinant progeny, plant #6. This revealed a fully intact recombination product, differing from the expected sequence by just one SNP within the CinH recognition site (Figure S7A).

**Figure 5:**
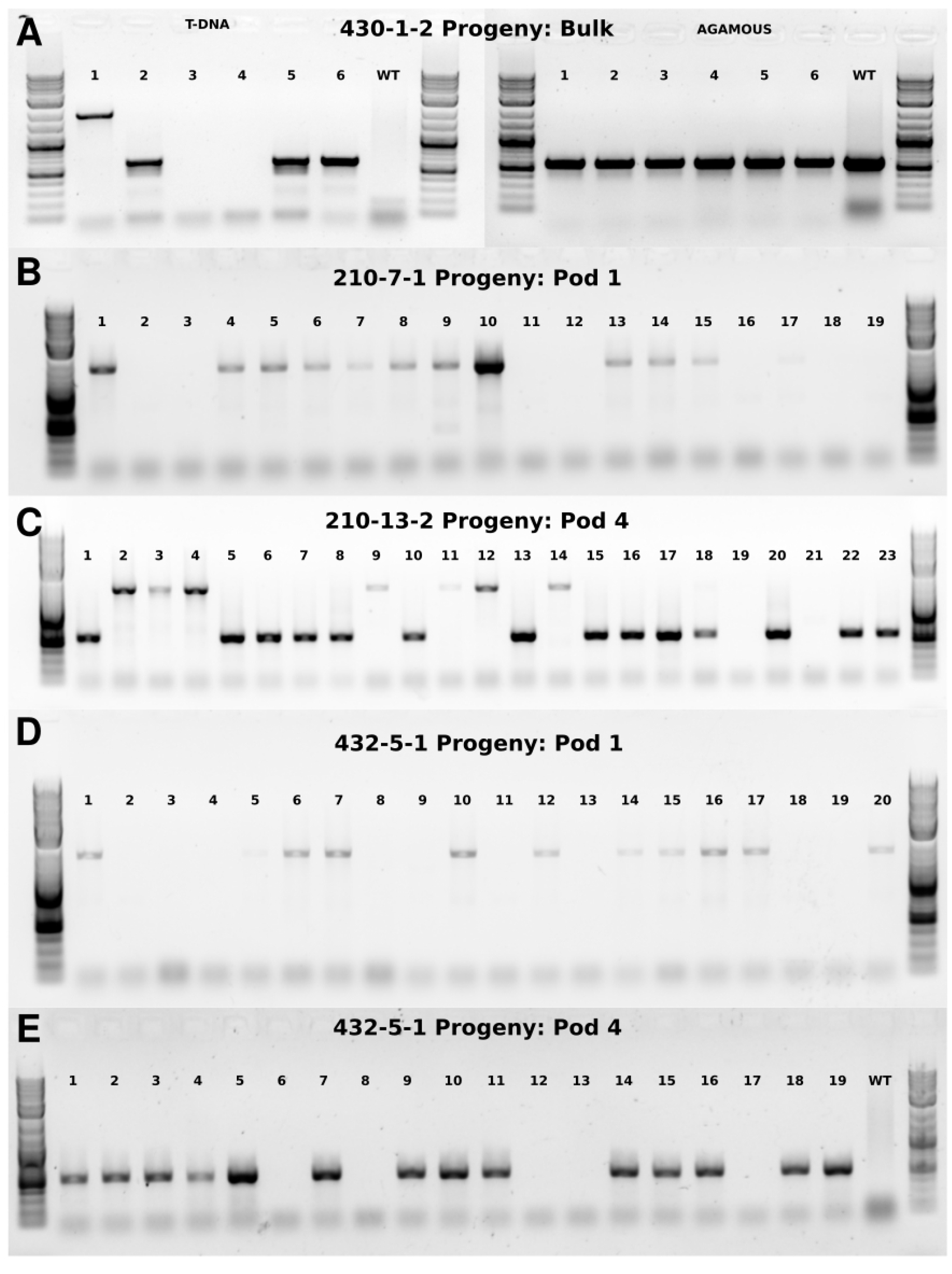
Heritable recombination events from CinH, FLP, and Integrase13. **5A:** Diagnostic PCRs for recombination and genomic control locus in 6 viable progeny of 430-1-2. *AGAMOUS* band is 594bp. **5B-C:** Diagnostic PCRs for recombination in pod-specific progeny of Integrase 13 plants 210-7-1 and 210-13-2. **5D-E:** Diagnostic PCRs for recombination in pod-specific progeny of FLP plant 432-5-1.

For Integrase13, we collected seed pod-by-pod from two of the infected parents: plants 210-7-1 and 210-13-2.

Selecting the first (lowest) pod from 210-7-1, we germinated and screened 19 progeny for recombination by PCR: 11 produced full-length bands, and 8 produced no band, indicating a heterozygous parent with no heritable recombination. Next, we selected the fourth pod from 210-13-2, germinating and screening 23 progeny: among these, 21 produced bands, including 7 full-length, and 14 recombination-length bands (Figure 5B-C). Given that the two empty lanes may have represented null segregants or PCR failure, we conservatively estimated the heritable recombination rate at ∼61%.

For FLP, we collected seed pod-by-pod from the high-expression plant, 432-5-1. From the lowest pod, we germinated 19 progeny and screened by PCR for the T-DNA (Figure 5D). Half the plants produced no band, indicating a heterozygous parent, and half produced the full-length band, indicating no heritable recombination. However, given that the infected parent plant showed much stronger Ruby signal in the upper portion of the plant (Figure 4C), we next germinated and screened an additional 19 progeny from the fourth pod up the plant. Many of these plants had red roots, and each of the 14 progeny yielding an amplicon for the T-DNA produced a recombination-length band, indicating a heritable recombination rate of at least 73% (Figure 5E).

## Discussion

Our work advances viral recombinase delivery in several respects. While this is not the first report of viral-mediated marker removal, the use of TRV vectors that efficiently access the meristem greatly simplifies recovery of recombinant progeny by bypassing the regeneration step typically required with other viral vectors. Advantages are also found in the expanded set of tested recombinases. Given wide-ranging results from comparative efficacy studies for SSRs in plant and mammalian systems^27–29^, it seems reasonable to expect some difference in recombination efficiency might be observed in our viral delivery system. However, all four tested SSRs appeared to saturate recombination within regions of stable infection. It could be that strong expression from TRV, or the proximity of recognition sites for the excision reaction, were sufficient to mask intrinsic differences that might be observable for more complex recombination reactions. With respect to viral vector stability, the larger recombinases FLP and Integrase13 did exhibit partial collapse on TRV. We nevertheless observed widespread Ruby activation, persisting through flowering, in multiple plants, indicating that all four vectors represent a viable means to achieve marker removal through infection. Beyond excision for marker removal, additional recombinases may also be useful for applications such as gene stacking^30–32^ and synthetic genetic circuits^33^, that rely on orthogonal activity of multiple SSRs.

All previous reports for viral-mediated marker removal used Cre, which we found frequently led to phenotypic penalties among infected plants, including reduced plant viability and fertility. Infections with Integrase13, FLP, and CinH vectors were similarly effective to Cre for inducing recombination, but had almost no impact on plant development and fertility. The only plants with reduced viability, from CinH event 430-1, could be due to early infection, as all plants from other events infected later with TRV-CinH were phenotypically normal. Alleviation of phenotypic defects with the other SSRs could be attributable to different and, for CinH and Int13, longer putative recognition sites that provide greater specificity in the context of the host genome. This is an important consideration for implementing viral recombinase delivery in other plant species, where new genome-specific context effects may arise.

Beyond direct applications of recombinases, our work prototypes a generalizable strategy for tracking viral infection with utility for development of future viral vectors. A primary challenge in most viral delivery systems is plant-to-plant variation in infection efficiency, compounded by the potential for vector collapse. The use of reporters to track movement is vastly preferable to RT-PCR, which is laborious, slow, and low-resolution. While some initial effort is required to develop and identify transgenics with sufficient expression, the excision-activated Ruby reporter offers significant advantages against multiple classical reporters: unlike luciferase or *β*-glucuronidase, it requires no substrate, and unlike fluorescent reporters, requires no specialized imaging equipment. Furthermore, in contrast to reporters encoded directly on the virus that reveal only active infection, the target line-embedded Ruby switch reveals entire infected cell lineages, including formerly infected cells and mitotic lineages of new growth. This is particularly relevant for tracking meristem invasion and germline modifications. For example, TRV meristem invasion is thought to be transient^34^, such that derived somatic and germline cells may carry recombination events in the absence of active infection. Previous VIGE studies have tracked “infection-derived” lineages by targeting native loci that produce phenotypes, such as PDS or magnesium cheletase, but chlorotic and bleaching phenotypes can be inconsistent and incur fitness penalties.

In this study, all four recombinase vectors were able to produce heritable modifications. Almost all the plants we screened using Ruby signal as an indicator for plant and pod selection produced heritable modifications, and in over 50% of the progeny. Notably, the only two pods that failed to yield recombinant progeny, from Integrase13 plant 210-7-1 and FLP plant 432-5-1, were the lowest (first) pods. This is consistent with our previous VIGE study with TRV, in which later-developing pods showed higher rates of heritable modification^3^. Previous work has also found variable effects on both somatic and germline editing efficiency from inclusion of mobile RNA signals such as the Arabidopsis Flowering Time (FT) motif, which may enhance mobility of viral subgenomic transcripts (including sgRNAs) or the viral genome itself to improve access to the meristem^3,35^. Our TRV recombinase vectors included no such signals, yet were able to reach germline cells, consistent with a model of direct meristem access via TRV.

In summary, we have demonstrated an effective approach for viral-mediated recombinase delivery, and developed a reporter platform for viral infection that may be applied for rapid optimization of vector architecture and delivery conditions in *N. benthamiana* and potentially other species.

## Materials & Methods

### Vector construction and strain preparation

T-DNA vectors for the *N. benthamiana* target lines were constructed via Golden Gate assembly, using a hybrid hierarchical approach to install individual components as Phytobrick parts^36^ into the modular system described in Cermak *et al*. (2017)^37^. Inserts for Phytobrick assembly were produced by PCR (ORFs, promoters, terminators) or as annealed oligos (recombinase recognition sites). The T-DNA vector with TRV1 (pNJB069) was previously described^3^. TRV2 vectors were constructed via Golden Gate assembly, using a TRV2 cloning backbone also previously described^3^. The inserts for Cre, enhanced FLP (FLPe), and Integrase13 were synthesized (TWIST Bioscience) following previously published sequences, lacking an NLS in each case. The CinH insert, including an SV40 NLS, was also synthesized, following a sequence kindly provided by James Thompson (USDA-ARS-CIG). All clones were propagated in NEB10*β* Competent cells (NEB C3019I), and confirmed via Sanger (Eurofins Genomics) or Oxford Nanopore (SNPSaurus) sequencing. Final T-DNA vectors were transformed into Agrobacterium strain GV3101, including the pMP90 helper plasmid.

### Generation of *N. benthamiana* target lines and agroinfection of TRV vectors

Transgenic *N. benthamiana* lines were produced following the method of Sparkes *et al*. (2006)^38^. TRV infections were initiated via leaf infiltration using, using TRV1 and each respective TRV2 vector together at a 1:1 ratio and an aggregate OD600 of 0.6.

### Genotyping of infected plants

Genomic DNA was collected from 1 cm leaf punches using a urea extraction protocol^39^. Diagnostic PCR was conducted using JumpStart REDTaq ReadyMix (Sigma Aldrich) with one of the following primer pairs for the T-DNA: CGCACAATCCCACTATCCTT and ACAACGATGGTGGTCACAGA, or CCACT-GACGTAAGGGATGACG and GATCGGAAGCGGCTTTGGT. The genomic control locus *AGAMOUS* was amplified with the following primer pair: CTTCCAGCCCTTTCTTTCCT and GAAAGGGGAGGAC-CTTTCAA. To simplify cloning and sequencing of representative amplicons, these primers were modified with Golden Gate extensions for assembly into MoClo destination vectors^40^. RNA was extracted from four 1 cm leaf punches, taken from different leaves, using the Direct-Zol RNA Miniprep kit (Zymo). RT-PCR was conducted using a OneStep RT-PCR kit (Qiagen) with the following primer pair: ACAACTCGGTTTGCTGACCT and ACCGATCAATCAAGATCAGTCGA.

## Supporting information

Supplementary_Fileset_1

## Supplementary

### Supplementary Figures

**Figure S1:**
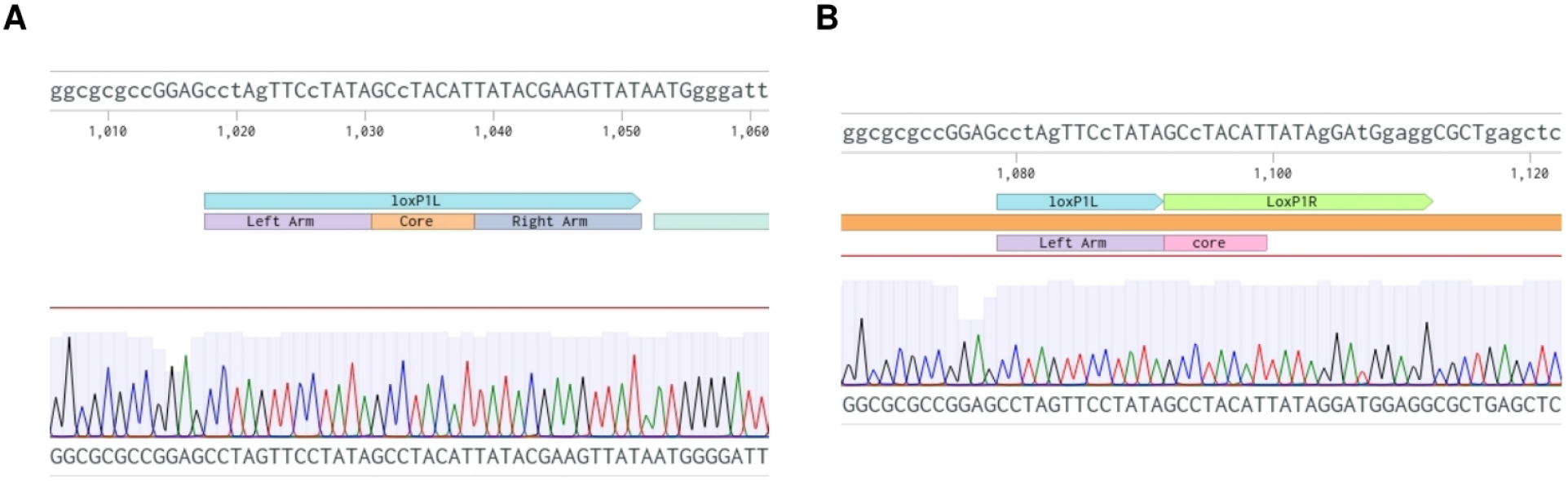
Sequencing of amplicons from diagnostic PCR in target line 263-8 infected with TRV-Cre. **1A:** Sanger read alignment of full-length band amplified from green tissue against the plasmid map for pJPC263. **1B:** Sanger read alignment of excision-length band amplified from red tissue against the predicted recombination product.

**Figure S2:**
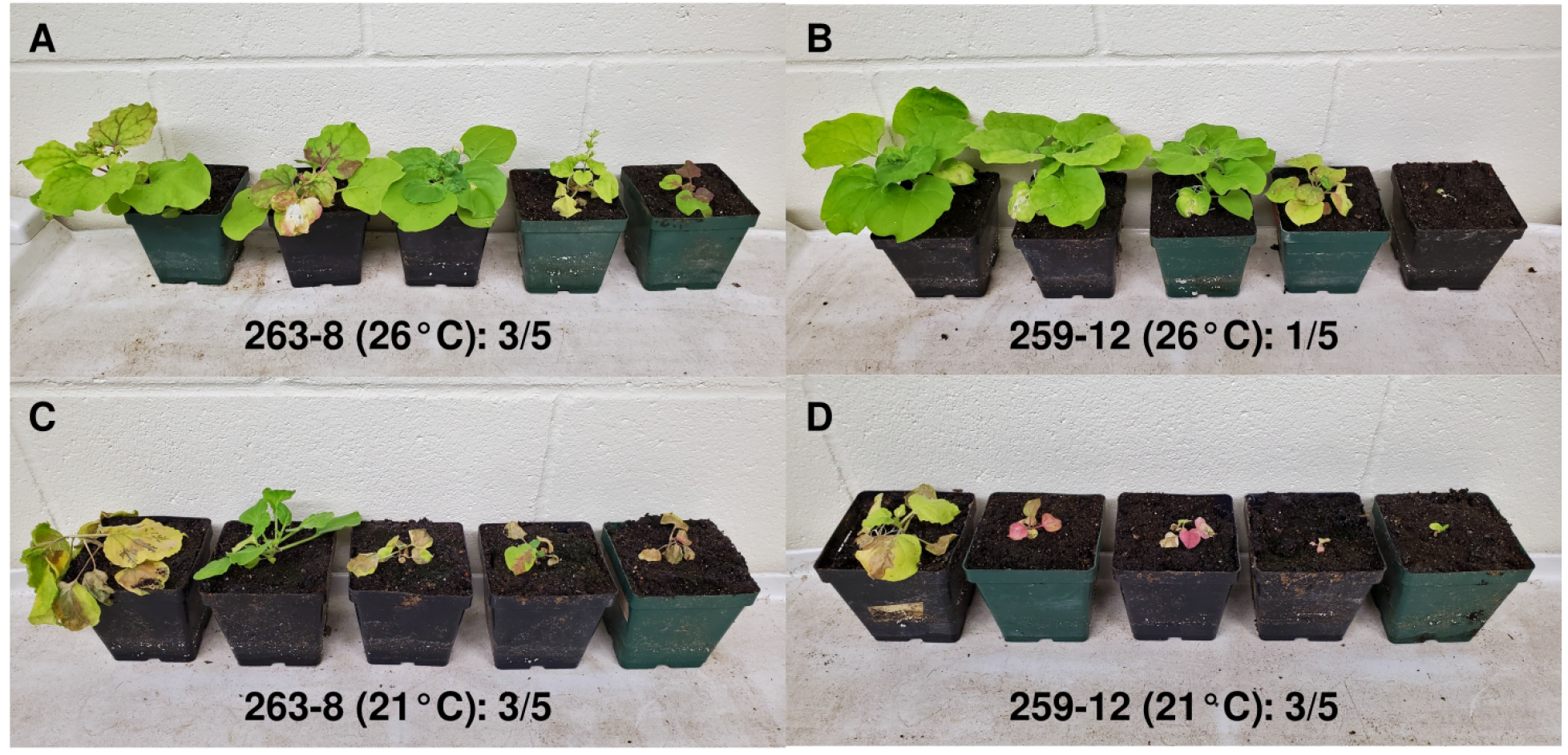
Ruby scoring and attrition among TRV-Cre infected plants. **S2A:** Plants from event 263-8 grown at 26°C. **S2B:** Plants from event 259-12 grown at 26°C. **S2C:** Plants from event 263-8 grown at 21°C. **S2D:** Plants from event 259-12 grown at 21°C.

**Figure S3:**
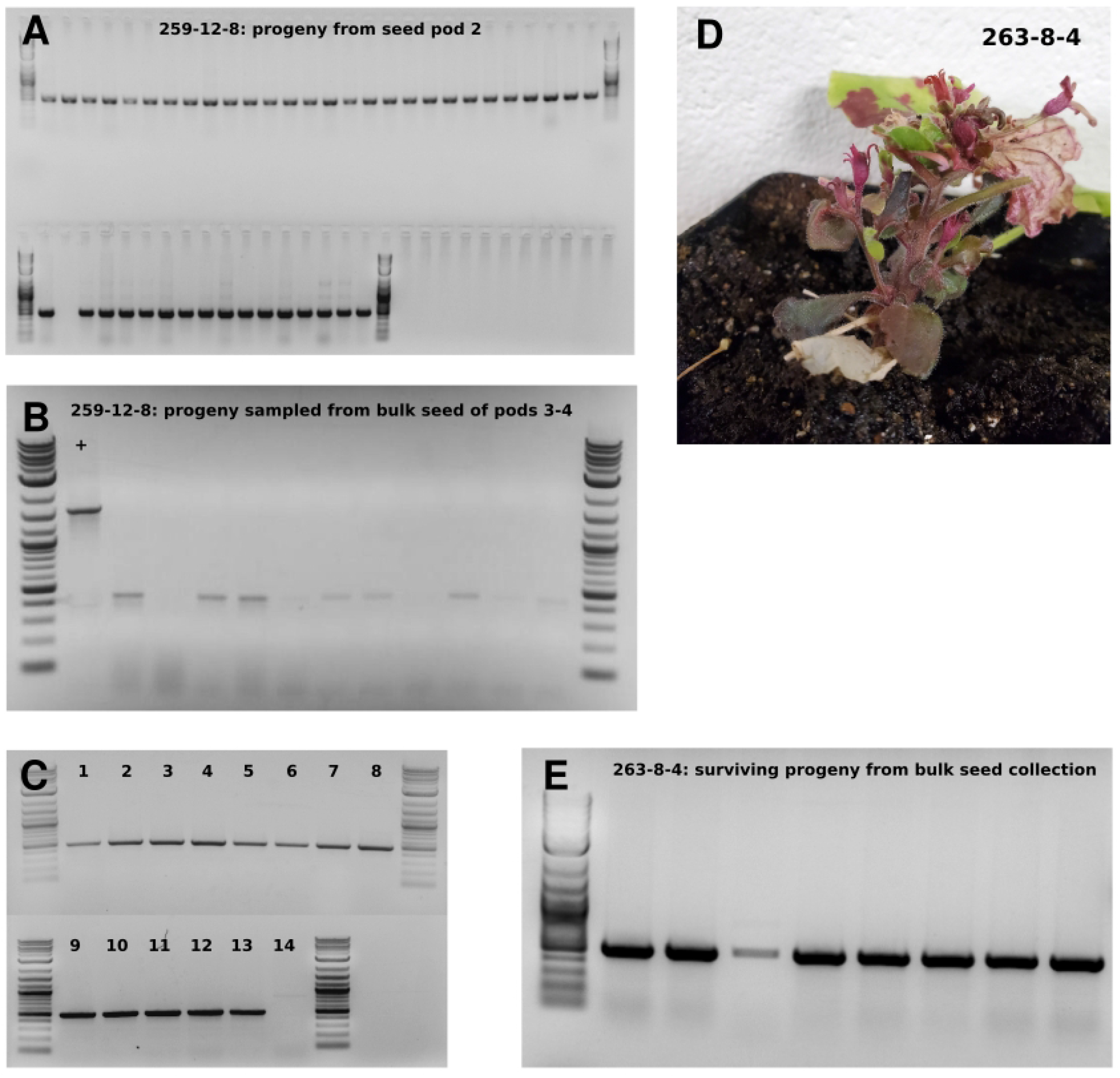
Genotyping additional progeny from TRV-Cre infected plants. **S3A:** Diagnostic PCR results from 46 progeny from seed pod 2 of 259-12-8. 45 out of 46 produced a positive band for recombination. **S3B:** Diagnostic PCR results from 11 progeny sampled from 25 total seed bulked from 259-12-8 pods 3 and 4: + indicates plasmid positive control. 7 out of 11 samples produced a clear positive band for recombination. **S3C:** Diagnostic PCR for subset of 14 seedlings from 3E. (Seedling 14 was fully green.) **S3D:** Plant 263-8-4 exhibited stunting, and produced few viable seed. From 52 total seed, 8 germinated, but died soon after. Genomic DNA was collected from these 8 seed and subjected to the diagnostic PCR. **S3E:** Diagnostic PCR results from 8x progeny of 263-8-4, all of which were positive for recombination.

**Figure S4:**
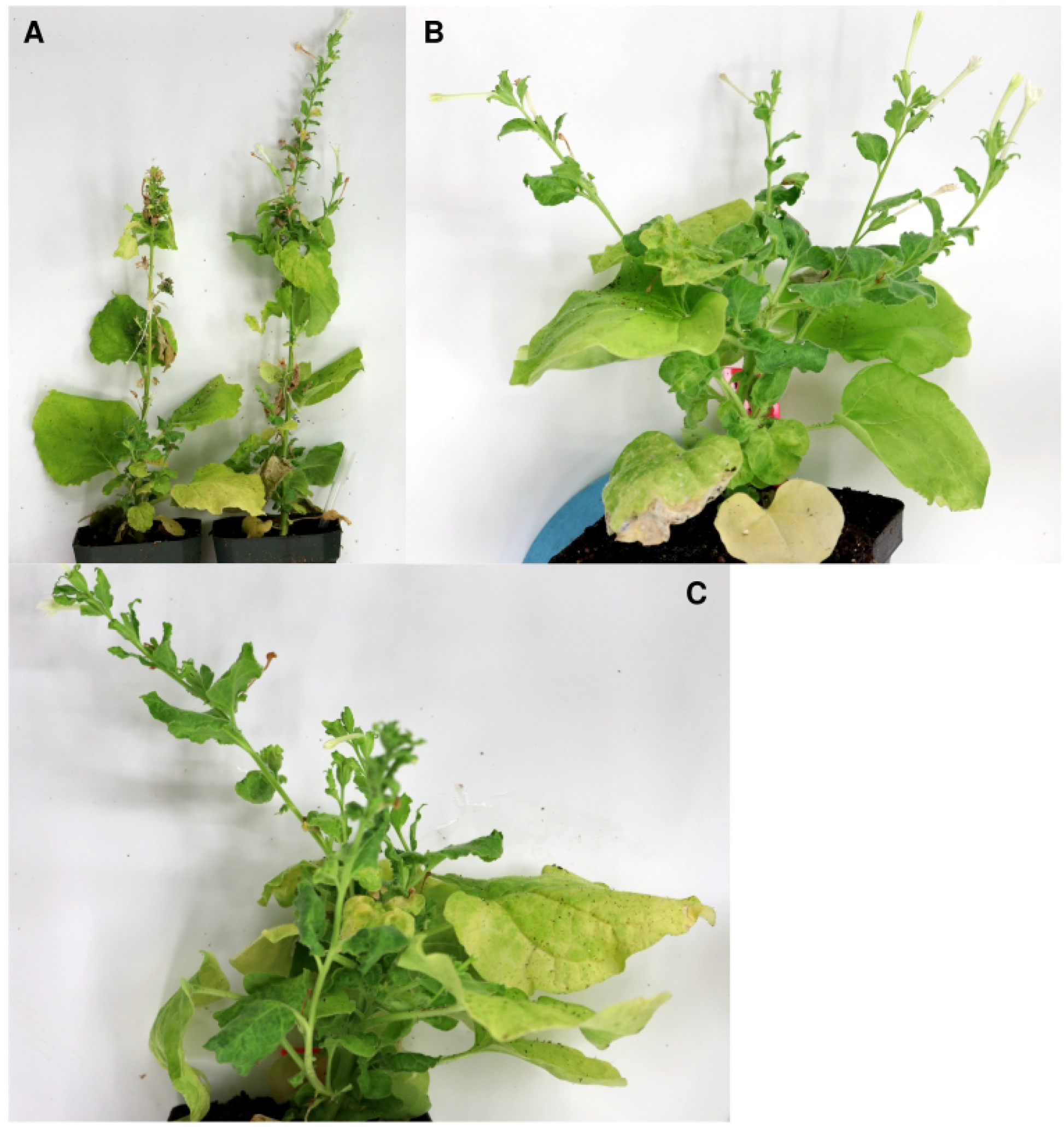
Phenotypic abnormalities among TRV-Cre infected plants. **S4A:** Wild-type plants infected with TRV-Cre showed phenotypic abnormalities, including stunting, misshapen leaves, and loss of apical dominance. Two of six infected plants are shown here. **S4B-C:** Putative transgenic target line plants infected with TRV-Cre but with no Ruby signal also showed defects, particularly loss of apical dominance.

**Figure S5:**
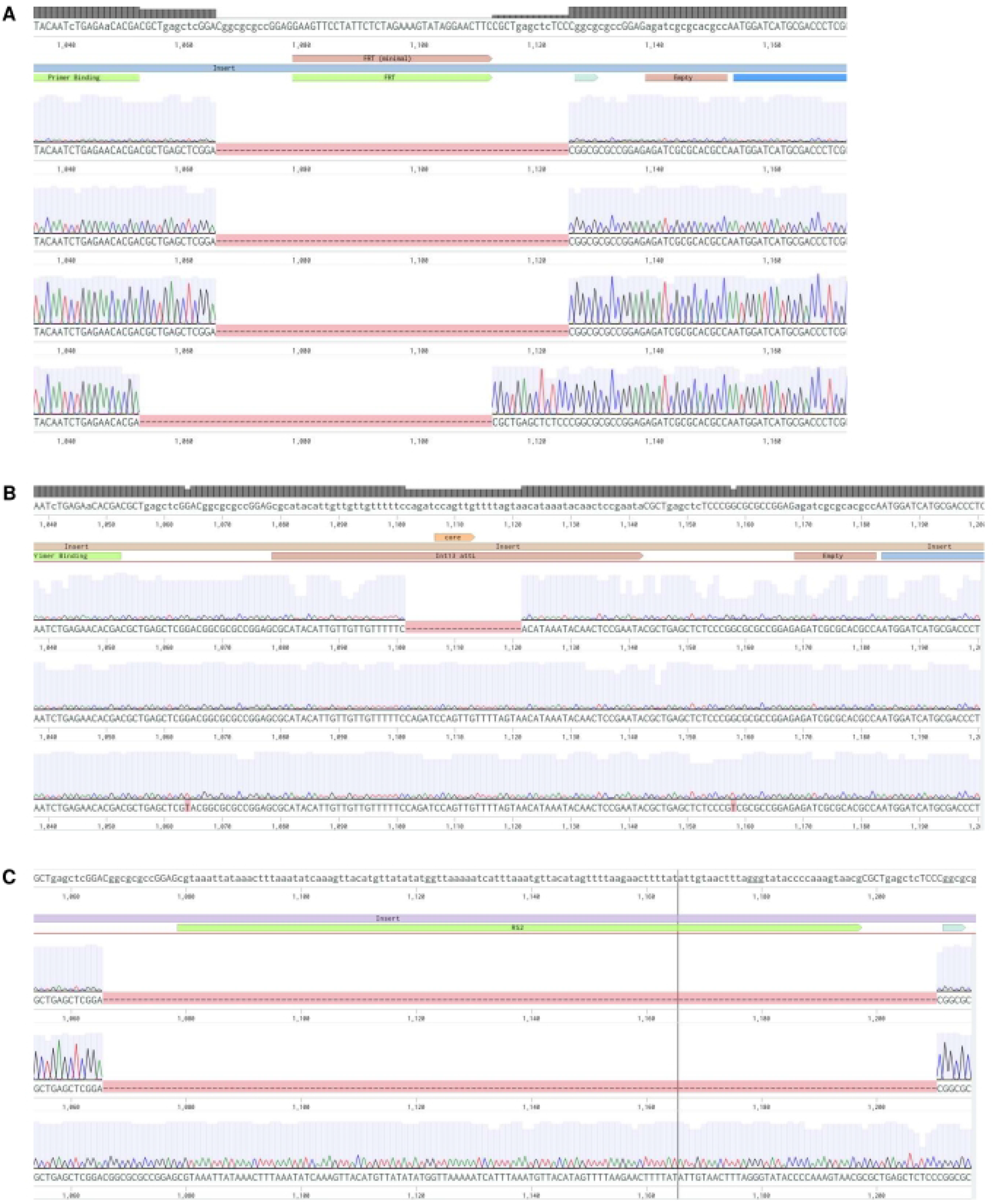
Sequencing clones of recombination bands for FLP, CinH, and Integrase13. **S5A:** Four clones from FLP. **S5B:** Three clones from Integrase13. **S5C:** Three clones from CinH.

**Figure S6:**
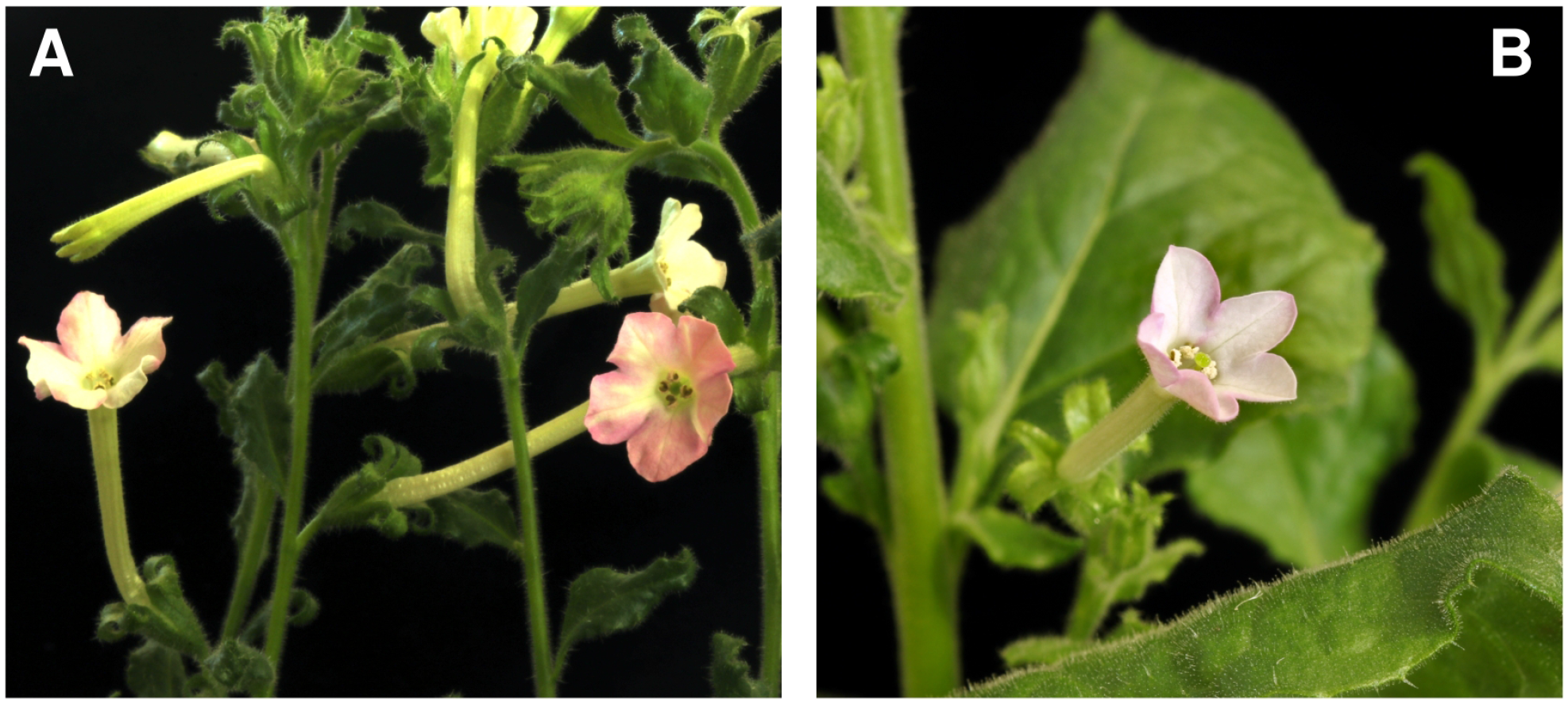
Ruby signal in flowers from additional Integrase13 plants. **S6A:** Plant 210-8-3. **S6B:** Plant 210-14-1.

**Figure S7:**
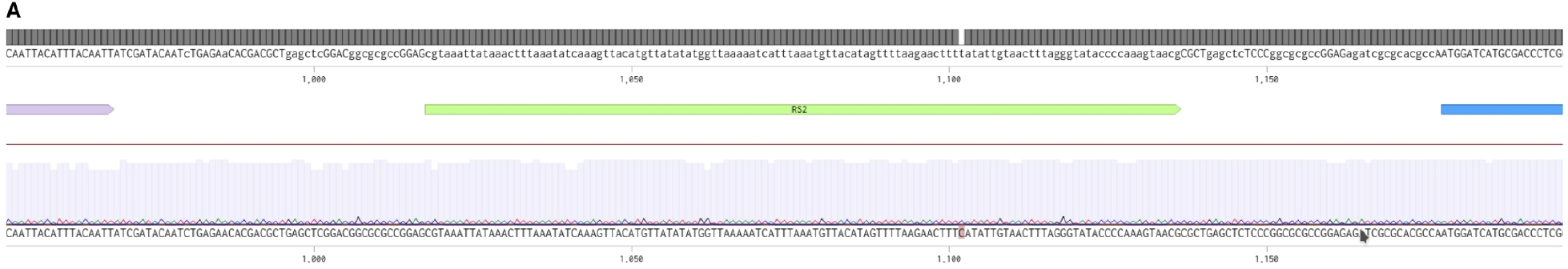
Sequencing clone of recombination bands from progeny of other TRV-recombinase plants. **S7A:** Plant 430-1-2-6.

### Supplementary Fileset 1

- Vector maps for constructs

